# A low-androgen environment sustains the functionally reduced sGC states required to restrain castration-resistant prostate cancer growth via elevated cGMP levels

**DOI:** 10.64898/2026.10.12.753798

**Authors:** Clara Troccoli, Beatriz Mateo-Victoriano, Annie Beuve, Merce Jorda, Janaki Sharma, Priyamvada Rai

## Abstract

Castration-resistant prostate cancer (CRPC) remains incurable and fatal, creating a dire need for new safe and effective treatment strategies. We recently reported that the soluble guanylyl cyclase (sGC) stimulator, riociguat, cooperates with standard-of-care androgen deprivation (AD) to inhibit castration-resistant tumor burden by oxygenating hypoxic tumor niches and inducing tumor apoptosis. We further found that AD restores the sensitivity of CRPC cells to riociguat, which requires a redox-active functional sGC complex to stimulate cyclic GMP (cGMP) production. As redox reprogramming is a feature of the low-androgen PC milieu, to better establish the parameters of riociguat efficacy, we wished to determine whether redox-inactive forms of sGC persist in the castrate setting. To do so, we treated CRPC LNAI tumor-bearing animals with cinaciguat, an sGC activator which preferentially acts on the oxidized sGC complex. We found that, compared to riociguat, cinaciguat did not significantly increase intratumoral or systemic cGMP levels in the castrate setting. Nor did cinaciguat treatment significantly decrease tumor burden or oxygenate the hypoxic CRPC tumors. Given the preferential activity of cinaciguat on the redox-inactive sGC complex, our results indicate that the low-androgen castrate setting maintains the sGC complex in its functionally reduced state, affirming the critical benefit of AD for riociguat efficacy in the lethal castration-resistant disease stage.

## INTRODUCTION

Prostate cancer (PC) is the most commonly diagnosed cancer among men worldwide (seer.cancer.gov) with five-year survival rates around 35% once the disease progresses to the metastatic castration-resistant (CRPC) stage^1^. Thus, there is a critical need for new, safe and effective treatments that improve the efficacy and durability of backbone androgen deprivation therapy (ADT)^2^. We recently reported that the clinical soluble guanylyl cyclase (sGC) agonist, riociguat^3^, inhibits CRPC growth in preclinical models in the castrate setting, with intratumoral and systemic elevation of its catalytic product cyclic GMP (cGMP) as a biomarker of on-target anti-tumor activity^4^. Elevated sGC activity improves tumor oxygenation and vascular structures in castration-resistant tumors, leading to tumor apoptosis and radiosensitization^4^. Thus, riociguat-induced stimulation of the sGC-cGMP axis eliminates hypoxic tumor niches that house aggressive PC cell populations associated with disease progression and treatment resistance.

The ability of riociguat to stimulate cGMP product is critically dependent on the presence, albeit even low levels, of the functional sGC obligate heterodimer^3,5,6^. The sGC complex is functionally regulated at level of protein oxidation, at its heme moiety and other residues^7-9^. Our prior study^4^ established that when androgens are present in culture, hormone-sensitive PC cells are significantly more responsive to riociguat than their castration-resistant counterparts. Riociguat is unable to stimulate cGMP production from the oxidized sGC complex^9^, suggesting that progression to castration resistance from the hormone-sensitive stage is associated with a shift to oxidized sGC^4^. Consistent with these observations, we found that androgen deprivation (AD) restores the ability of riociguat to elevate cGMP levels and suppress in vivo castration-resistant PC growth^4^. This observation strongly suggests that androgen blockade, a standard-of-care treatment for advanced PC, functionally regenerates the reduced sGC complex. In our preclinical in vivo studies^4^, riociguat significantly decreased CRPC burden compared to control cohorts. However, riociguat did not achieve equivalent tumor suppression across all animals in the cohort, with the extent of anti-tumor effect correlating positively with intratumoral and systemic cGMP levels. Thus, we wished to determine whether non-responsive tumors represent the persistence of oxidized sGC states in the in vivo castrate setting, as these would negatively impact riociguat efficacy in stimulating cGMP production. This aspect is especially critical to elucidate as the low availability of androgen ligands produced by AD alter the redox milieu and androgen receptor (AR) function^10-13^.

Although establishing the sGC redox state is critical to the effective use of different sGC agonists, it is difficult to directly detect oxidation states of the complex, especially in vivo. Therefore, in these studies we utilized the sGC activator cinaciguat (BAY58-2667) as a specific probe for the oxidation state of sGC in the in vivo castrate setting, as cinaciguat preferentially increases cGMP production from the oxidized sGC complex^14,15^. This selectivity of cinaciguat has previously been reliably leveraged as a surrogate marker for sGC oxidation^9,16^. Using the same preclinical LNAI CRPC model as in our riociguat study^4^, we found that cinaciguat treatment could not suppress CRPC growth nor elevate intratumoral or systemic cGMP levels capable of oxygenating CRPC tumors. The relative inefficacy of cinaciguat compared to riociguat indicates that sGC exists predominantly in the functionally reduced state in the castrate setting, affirming the importance of androgen blockade-induced redox reprogramming^11^ for reductively regenerating the sGC complex in the lethal CRPC disease stage.

## RESULTS

### Cinaciguat treatment does not significantly decrease CRPC tumor growth and burden

Our prior studies have shown that PC cell lines exhibit hormonal status-specific sensitivities to sGC agonists. Specifically, hormone-sensitive cell lines show a marked sensitivity to riociguat under both androgen-replete and androgen-deprived conditions, whereas progressively more castration-resistant counterparts show either complete resistance to sGC agonists due to loss of the functional sGC heterodimer or a preference for cinaciguat under androgen-replete conditions^4^. Under the clinically relevant low-androgen castrate conditions, advanced CRPC cells show improved response to riociguat both in vivo and in vitro^4^, indicating a shift from the oxidized to functionally reduced sGC complex. However, the relatively high micromolar IC_50_ for riociguat even in the responsive cell lines indicates the persistence of oxidizing conditions in vitro, making it difficult to assess the full extent to which AD regenerates the functionally reduced sGC complex.

To determine whether oxidized sGC states persist under in vivo AD and thus compromise riociguat efficacy, we tested whether cinaciguat can restrain in CRPC growth in a similar xenograft tumor setting as our prior riociguat study^4^. To this end, we subcutaneously injected 2x10^6^ LNAI CRPC cells into the flanks of 6 week old Nu/Nu castrated male mice, which lack adrenal androgens^17^, to recapitulate androgen deprivation. Following palpable tumor growth, animals were randomized into groups that received intraperitoneal (IP) injections of either vehicle (DMSO) or 20 mg/kg/day cinaciguat daily. This cinaciguat dose had no adverse systemic effects on the study animals (**Fig. 1A**). Tumor growth rates were monitored using caliper-based measurements of the subcutaneous tumors. Unlike riociguat^4^, cinaciguat treatment did not significantly affect overall tumor kinetics or endpoint tumor weights/volumes relative to the vehicle groups (**Figs. 1B-E**). Cinaciguat treatment also did not impair the ability of prostate-specific antigen (PSA) to predict castration-resistant tumor burden (**Fig. 1F**), an important consideration when evaluating combinatorial strategies with backbone AD for translation into the clinic.

**Figure 1.**
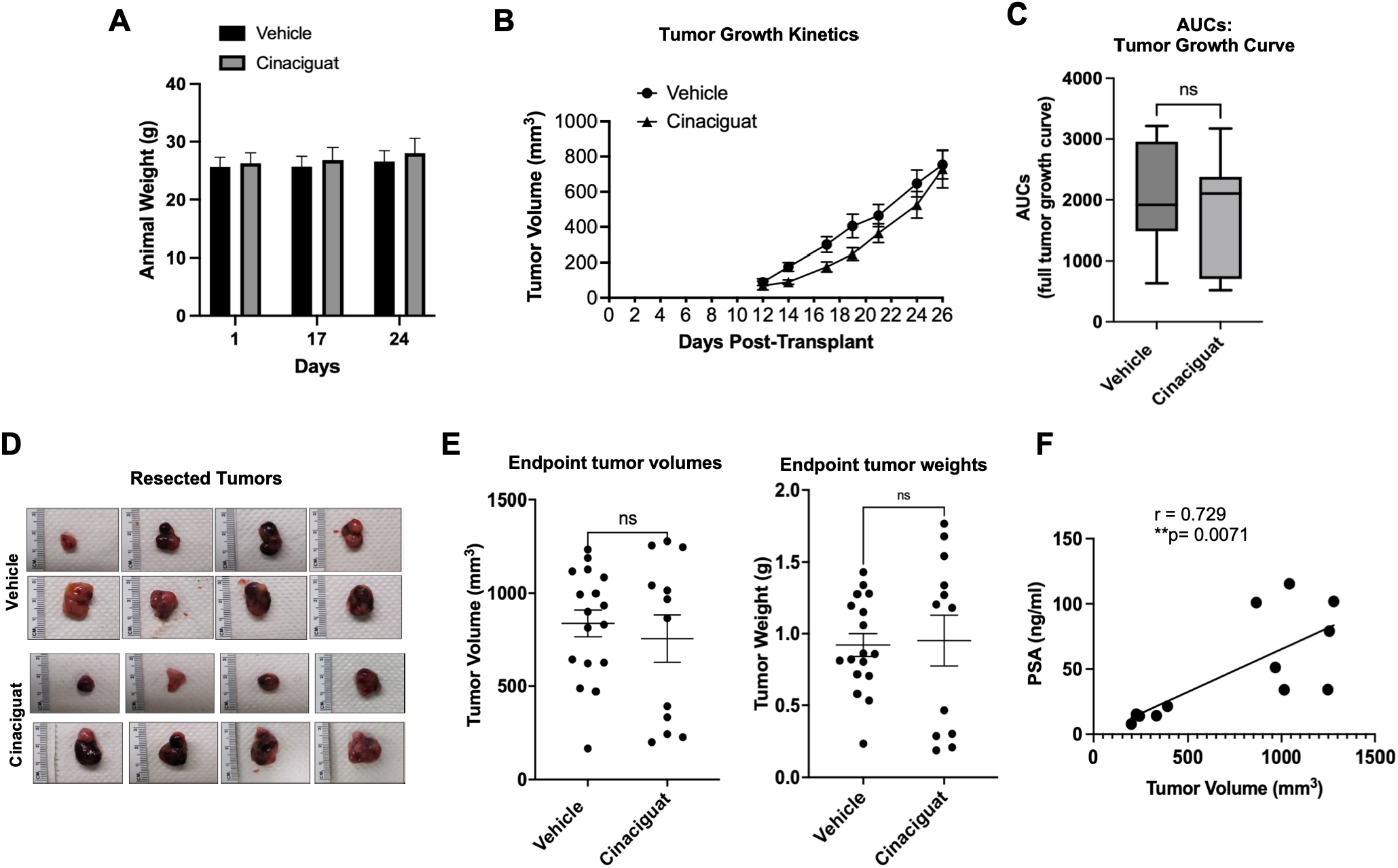
Cinaciguat treatment does not significantly inhibit CRPC growth. A) Animal body weights in the two treatment cohorts. Averaged weights with standard error of the mean (SEM) are shown. B) Tumor growth kinetics for LNAI tumor-bearing castrated male Nu/Nu mice from cinaciguat (20 mg/kg/day) or DMSO treatment groups Averaged volumes for the indicated timepoints with SEM are plotted. C) Averaged Area-Under-Curve (aAUC) values for tumor growth. Unpaired two-tailed Student’s t-test, with Welch’s correction. D) Representative images of castration-resistant tumors derived from the two treatment groups. Ruler shown for scale to the left of the tumors. E) Endpoint tumor volumes (mm^3^) and tumor weights (g) per treatment group. Each point represents an individual tumor. Averaged values with SEM are shown. Unpaired two-tailed Student’s t-test, with Welch’s correction. F) Correlation between circulating PSA levels vs. tumor volumes in the cinaciguat-treated group. Pearson correlation coefficient, r, and the significance of the correlation, p, are shown.

### The low-androgen redox milieu induces intratumoral and systemic sGC states that are refractory to cinaciguat-mediated stimulation of cGMP production

In our prior study^4^, we found that inhibition of castration-resistant LNAI tumor growth correlated significantly with riociguat-mediated increase in intratumoral as well as circulating levels of cGMP, reflecting on-target stimulation of sGC catalytic activity. Given the lack of overall anti-tumor efficacy from cinaciguat treatment (**Fig. 1**), we next assessed whether its lack of efficacy arose from an inability to stimulate cGMP production in the CRPC setting. To this end, we measured intratumoral and systemic cGMP in vehicle vs. cinaciguat-treated animals and compared them to levels in archived tissue and sera collected from riociguat-treated animals in our prior study^4^.

Unlike riociguat, cinaciguat treatment did not significantly elevate intratumoral cGMP levels (**Fig. 2A**). To determine more directly whether the relative unresponsiveness of LNAI tumors to cinaciguat vs. riociguat reflected their specificities for different sGC redox states, we moved three animals that had been on cinaciguat treatment for approximately 3 weeks, to riociguat for 72 hours. At the end of this time period, we euthanized these animals and compared their intratumoral cGMP levels to three animals who had remained on cinaciguat, controlling for tumor size. Despite the short duration of this switch at a timepoint when the tumors were fairly large (**Fig. 2B**, left) there was a noticeable albeit non-significant trend of elevated intratumoral cGMP following riociguat treatment relative to the tumors that remained on cinaciguat (**Fig. 2B**, right). Nevertheless, these results are consistent with a predominantly reduced sGC state in the castrate setting.

**Figure 2.**
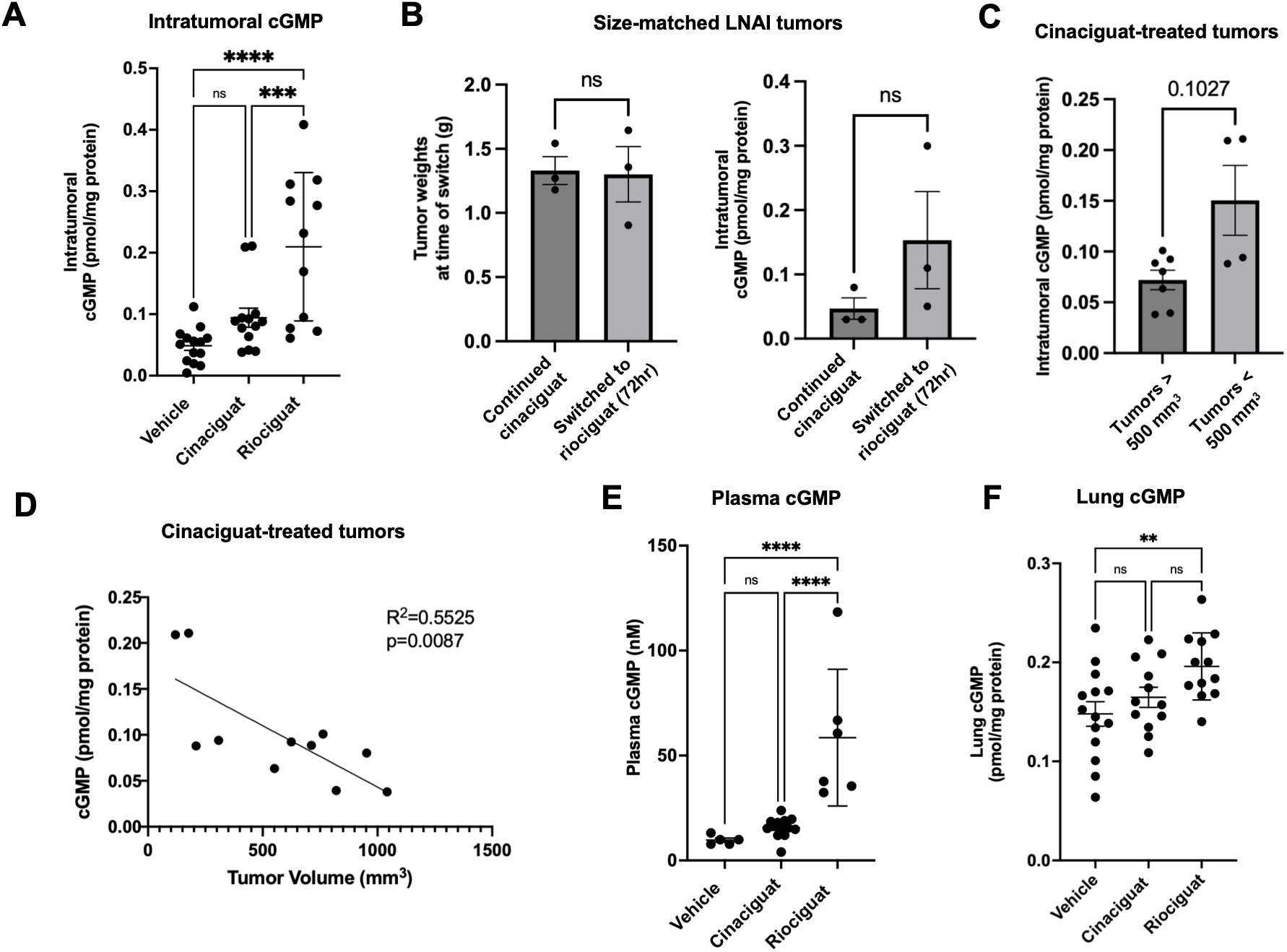
Cinaciguat does not significantly stimulate cGMP levels in the castrate setting. A) Intratumoral cGMP levels. Tumor tissue levels in the vehicle and cinaciguat-treated animals were compared with levels measured in archived tumor tissues from riociguat-treated animals. Each point represents an individual tumor. Averaged values with SEM are plotted. Ordinary one-way ANOVA with Fisher’s LSD test. B) Intratumoral cGMP levels from three animals who were switched following 3 weeks of cinaciguat treatment to riociguat (20 mg/kg) for 72 hours. Levels were compared to three animals who remained on cinaciguat with size-matched tumors. Left: Tumor weights for the comparison cohorts. Right: Averaged intratumoral cGMP values with SEM are shown. All p-values are from unpaired two-tailed Student’s t-tests, with Welch’s correction. C) Difference in intratumoral cGMP levels between cinaciguat-treated tumors larger than 500 mm^3^ vs. smaller than 500 mm^3^. Unpaired two-tailed Student’s t-test, with Welch’s correction. D) Correlation between cinaciguat-treated tumor volumes and intratumoral cGMP levels. Pearson correlation coefficient, p-value are shown within the graph. E) Circulating cGMP levels. Plasma levels from vehicle and cinaciguat-treated cohorts were compared with cGMP levels in archived plasma from riociguat-treated animals^1^. Each point represents an individual animal. Averaged values with SEM are plotted. Ordinary one-way ANOVA with Fisher’s LSD test. F) Lung cGMP levels. Lung tissue levels from the vehicle and cinaciguat-treated cohorts were compared with levels in archived lung tissue from riociguat-treated animals^1^. Each point represents tissue from an individual animal. Averaged values with SEM are plotted. Ordinary one-way ANOVA with Fisher’s LSD test.

To evaluate whether the suboptimal efficacy of cinaciguat was due to compromised drug potency or delivery, we then separately analyzed intratumoral cGMP levels in tumors from cinaciguat-treated animals that were larger than 500 mm^3^ (‘nonresponders’) or smaller than 500 mm^3^ at endpoint (‘responders’). The larger tumors had lower cGMP levels than the smaller tumors (**Fig. 2C**). Although the relative differences in cGMP levels fell just short of statistical significance, intratumoral cGMP levels showed a significant negative correlation with tumor volume (**Fig. 2D**), similar to what we had observed with riociguat^4^. These results suggest that, although cinaciguat is less efficacious than riociguat in stimulating cGMP production, the limited stimulation observed in the CRPC tumors correlated with a lower tumor burden as expected. Collectively, these data support our conclusion that the observed inability of cinaciguat to sufficiently stimulate cGMP in the CRPC setting is not the result of suboptimal drug delivery or potency but rather reflects the predominance of the functionally reduced sGC complex.

We next evaluated the systemic effects of cinaciguat treatment. Compared to riociguat, cinaciguat did not significantly elevate cGMP levels in plasma (**Fig. 2E)** or in lung tissue (**Fig. 2F**) relative to vehicle-treated cohorts. Given that lung is the physiologic site of sGC activity, this finding supports cinaciguat’s specificity for the oxidized sGC complex, as castrate conditions are not anticipated to induce oxidative stress in lung tissue from otherwise healthy animals. Androgen blockade has been reported to decrease inflammation in lung tissue^18,19^. Indeed, the inability of cinaciguat to stimulate cGMP levels either intratumorally or systemically in the castrate setting further confirms that the low-androgen milieu supports reductive regeneration of the sGC complex.

### Cinaciguat treatment does not induce cell death or oxygenation in LNAI CRPC

We next assessed whether cinaciguat treatment was associated with any tumor-inhibitory response. Blinded histopathologic scoring of vehicle vs. cinaciguat-treated tumor sections showed that there was no significant difference in either staining of the pan-proliferation marker Ki67 (>90% in all evaluated sections) or in markers of apoptosis or necrosis (**Figs. 3A, B**), consistent with the lack of significant difference in CRPC burden (**Fig. 1**). Riociguat treatment induces significant tumor oxygenation of the hypoxic CRPC tumors, consistent with the physiologic outcomes from sGC-cGMP stimulation^4^. We similarly evaluated tumor hypoxia in cinaciguat vs. vehicle tumors through staining with pimonadizole (Hypoxyprobe), which shows maximal fluorescence at pO_2_ < 10mm Hg^20^. Similar levels of fluorescent signal were detected in vehicle and cinaciguat-treated tumors (**Figs. 3C, D**), indicating cinaciguat is not as effective at oxygenating CRPC tumors as riociguat, consistent with the inability of cinaciguat to stimulate cGMP production in the castrate setting. Collectively, these data support that cinaciguat is unable evoke tumor-inhibitory phenomena required to restrain CRPC growth, further linking the ability to elevate cGMP levels from the functionally reduced sGC complex to suppression of CRPC.

**Figure 3.**
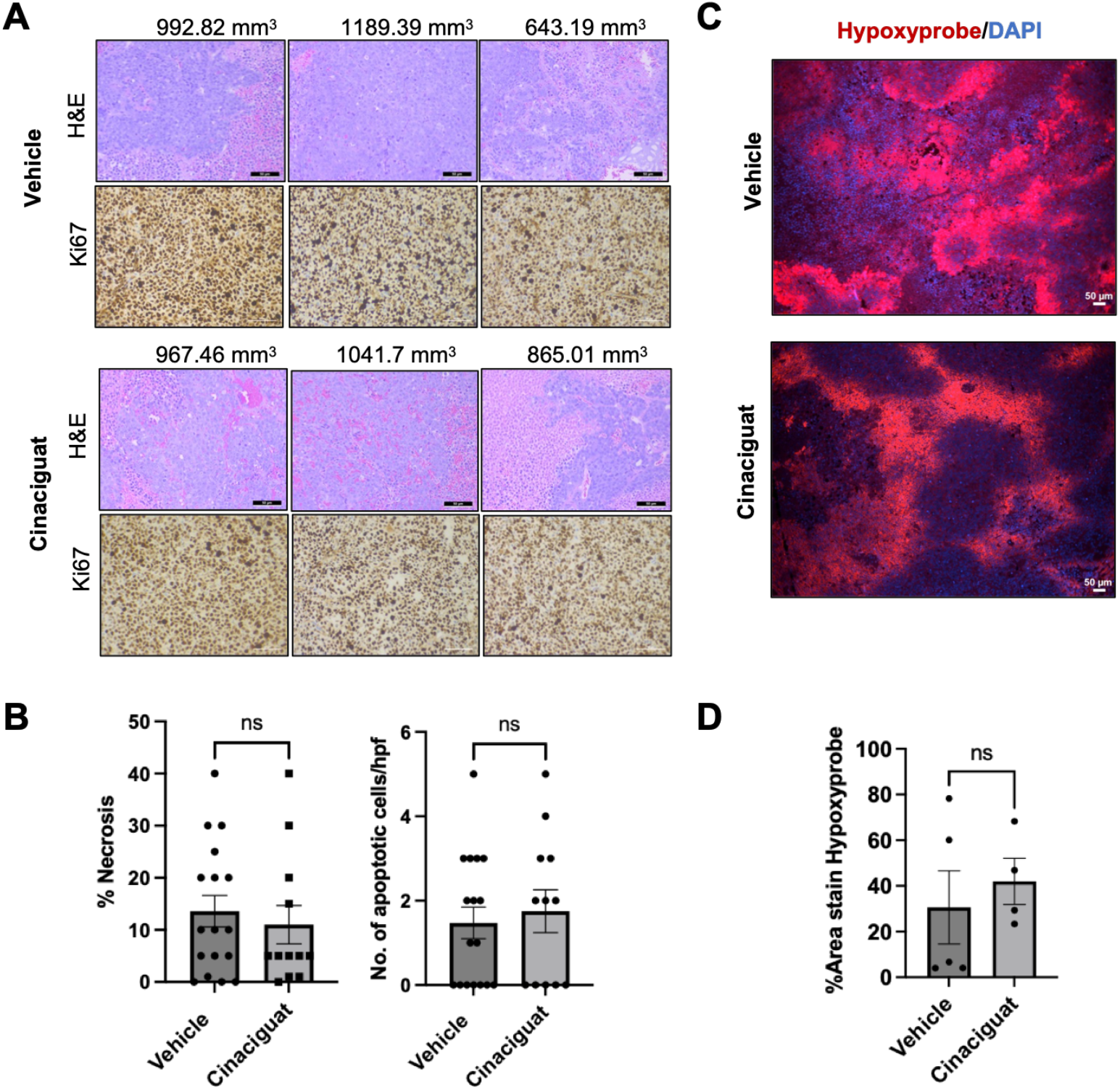
Cinaciguat does not induce cell death or tumor oxygenation in LNAI CRPC. A) Representative H&E and Ki67-stained images (50 μM scale bar) from vehicle and cinaciguat-treated tumors. B) Histopathological scoring for %necrosis (left) and number of apoptotic cells per high-power field (right) from tumor sections. Sections were scored blinded by a genitourinary cancer-specializing pathologist. Average scores with SEM are shown, with individual points representing distinct tumors. Unpaired two-tailed Student’s t-test, with Welch’s correction. C) Representative immunofluorescent Hypoxyprobe images (50 μm scale bar) from vehicle and cinaciguat-treated tumors. D) Quantitation of positive Hypoxyprobe-stained areas from a minimum of 4 tumors/treatment group. Each point represents averaged quantitation values from a minimum of 4 fields per tumor, with SEM. Unpaired two-tailed Student’s t-test, with Welch’s correction.

## DISCUSSION

Dynamic reprogramming of redox metabolism is a feature of many cancers, including prostate cancer^11,21^. Successful clinical repurposing of sGC agonists depends critically on establishing the physiologic sGC oxidation state, a key determinant of the differential efficacy observed in stimulators versus activators^22^. Our previous study^4^ demonstrated that the clinical sGC stimulator riociguat cooperates with AD to suppress tumors through enhanced cGMP production. These studies further found that riociguat efficacy in CRPC could be limited by methylation-mediated loss of the functional heterodimer or by oxidation of the sGC complex. Here, we used the sGC activator cinaciguat to determine the sGC oxidation state in the same in vivo LNAI CRPC model and to evaluate concomitant effects on its ability to stimulate tumor-inhibitory cGMP levels.

Our studies definitively show that cinaciguat, which preferentially stimulates cGMP production from the oxidized sGC complex^15^, is unable to suppress CRPC growth or significantly increase either intratumoral, circulating, or lung cGMP in the castrate setting. These results indicate that the low-androgen castrate milieu maintains the functionally reduced sGC complex not only in tumor tissue but also in non-tumor-bearing organs. This systemic effect is consistent with reports that androgen blockade exerts anti-inflammatory effects^18,19^. Importantly, these findings indicate that although AD-induced redox-protective reprogramming is a critical adaptation against oxidative stress associated with AR reactivation in CRPC^11^, this selfsame adaptation directly contributes to the therapeutic efficacy of riociguat by enabling sGC stimulation of cGMP production^4^.

Hypoxia is a key factor in driving metastatic castration-resistant prostate cancer growth and promoting resistance to standard-of-care treatments, including radiotherapy and androgen deprivation^23-28^. In our prior study, we found that riociguat treatment effectively oxygenates xenograft castration-resistant tumors, even those that remain relatively large following treatment^4^. This observation suggests that a lower threshold of cGMP stimulation is required for canonical sGC vasodilatory function relative to its anti-tumor efficacy. Nevertheless, tumor oxygenation induced by riociguat treatment has a radiosensitizing effect^4^. By contrast, the inability of cinaciguat to improve tumor oxygenation or induce tumor-suppressive responses supports that the elimination of hypoxic niches and apoptosis in riociguat-treated tumors^4^ is a direct effect of on-target sGC stimulation and elevation of cGMP by riociguat.

While our results underscore the lack of anti-tumor effects from cinaciguat treatment in CRPC, these studies do not necessarily preclude potential benefit from sGC activators in the broader cancer spectrum. Moreover, as the role of sGC-cGMP signaling in cancer expands, additional preclinical studies with sGC activators and stimulators are warranted to comprehensively evaluate the determinants of their therapeutic efficacy. Importantly, for the CRPC stage, AD is the standard-of-care and our results definitively demonstrate that the low-androgen milieu sustains the functionally-reduced sGC complex, as evidenced by the inefficacy of cinaciguat to stimulate cGMP production in this setting. Furthermore, by directly testing cinaciguat in the same in vivo CRPC model as our prior riociguat study, our findings strengthen the causal link between increased cGMP levels and tumor-suppressive responses, including enhanced tumor oxygenation. Thus, by ruling out significant representation of oxidized sGC states in the low-androgen castrate setting, this work affirms the translational potential of repurposing riociguat alongside backbone AD in metastatic CRPC, with cGMP levels serving as a biomarker for on-target treatment efficacy. More broadly, our study extends the concept of redox reprogramming in cancer beyond oxidative stress by demonstrating that the redox state of a clinically actionable signaling protein can be regulated by the hormonal environment, a critical consideration for cancers that rely on hormone therapy as a backbone treatment.

## MATERIALS AND METHODS

### Cell lines

LNAI CRPC cells were validated by short term-repeat (STR) profiling (Genetica Corp) and certified pathogen free prior to use. All cell culture reagents described in this section were obtained from Gibco, Life Technologies, except for sera, FBS and charcoal stripped serum (CSS), which were obtained from Hyclone. LNAI cells were cultured in RPMI-1640 medium (Gibco, 11875093). Media were supplemented with 5% FBS (Hyclone, SH30396) and 100 U ml^−1^ penicillin/streptomycin (Gibco, 15140122) for cell line growth and maintenance. Cells were cultured at 37 °C in a humidified incubator at 21% oxygen/5% CO_2_ or 5% O_2_/5% CO_2_ (HeraCell Tri-Gas, ThermoFisher Inc).

### Measurement of cGMP levels

Steady state cGMP concentrations in mouse plasma, tissues and human serum were measured by cGMP Enzyme Immunoassay (Arbor Assays, K065-H). Lysates were prepared according to the manufacturer’s instructions with minor modifications. Briefly, lung and tumor samples were pulverized in liquid nitrogen, 500 µl of sample diluent were added for every 50 mg of powdered lung tissue and 250 µl of sample diluent were added for every 50 mg of powdered tumor tissue. All lysates were then sonicated and incubated on ice for 10 minutes for uniform lysis, followed by centrifugation at 13,000 rpm for 10 minutes at 4°C. The BCA assay was used to measure protein concentration of the lysates and cGMP levels were expressed as picomoles of cGMP per milligram of total protein for tissue samples. Measurements in sera and plasma samples were carried out as follows. Plasma samples were diluted 1:5 in sample diluent and the cGMP signal read and calculated at 450 nm on a Spectramax iD3 or iDx Microplate Reader. All samples were run in triplicate. Final values were obtained by multiplying averaged plate reader values by dilution factor.

### Xenograft tumor experiments

All animal studies were performed in accordance with the University of Miami Institutional Animal Care and Use Committee (IACUC)-approved protocol. Numbers of animals to be used for the tumor-formation experiments were determined through power analysis to provide 90% statistical power to detect a mean difference of 2.6 between two groups, assuming a two-sample Student’s t-test and a standard deviation for both groups of 1.5 at two-sided 5% significance level.

For all xenograft experiments, cells were resuspended in a 1:1 matrigel (BD Biosciences, 356237): full FBS/RPMI-1640 media mixture and injected subcutaneously using a 26-gauge needle into one flank of immunocompromised 5–6-week-old castrated male mice (Nu/Nu, Envigo). Tumor length, width, and height were measured using electronic precision calipers (VWR, 90028). Tumor volumes were calculated according to the following formula: 0.52x(height*width*length).

For assessing whether cinacguat could mitigate in vivo CRPC tumor growth, we subcutaneously injected 2 × 10^6^ LNAI cells. Animals were randomized into a vehicle or treatment groups once tumors were palpable (100–150 mm^3^) and treatment was initiated with either DMSO:Tween80:PBS (10%:10%:80%) or 20 mg kg^−1^ cinaciguat (Cayman Chemicals, 17468) diluted in Tween80 and Mg^2+^ and Ca^2+^-free DPBS. Injections were given intraperitoneally (IP) seven days a week starting at palpable tumor formation.

For all xenograft experiments, tumor measurements and animal weights were monitored three times a week in a non-blinded manner. Tumor-bearing animals were euthanized when tumors in any group exceeded 10% of animal body weight (∼1000 mm^3^). Immediately following euthanasia, blood was collected through cardiac puncture from experimental animals and processed for serum and plasma. Tumors were excised and sectioned into samples for formalin fixation or snap-frozen in liquid nitrogen. Lungs were also collected and snap frozen for measurement of cGMP levels.

### Immunohistological staining

For histopathological analysis, fine sections (4 μm) were cut from formalin-fixed, paraffin-wax-embedded samples, and stained with hematoxylin and eosin. Immunohistochemical analyses were performed utilizing ready-to-use Ki67-specific antibody solution (K2, Leica Biosystems, PA0230). Sections were dewaxed, rehydrated, and pretreated in a high pH (pH 9) solution at 100 °C. Slides were dehydrated using a standard protocol and mounted using Permount. Ki67 images were taken using a Leica microscope and associated LAS V4.9 software at 40X objective magnification. Stained sections were scored blinded for histological scores by a pathologist.

### Immunofluorescent staining

For determination of hypoxia, pimonadizole (Hypoxyprobe, HP10-100) was diluted in saline and administered to study animals intraperitoneally at a dose of 60 mg/kg two hours prior to euthanasia. Tissues (tumor, lung, and liver) were then collected and fixed in formalin and paraffin embedded. Paraffin-embedded fine sections were deparaffinized by heating at 56°C for two hours and then hydrated by treating with Xylene (15 minutes, 2 times), 100% ethanol, 90% ethanol, and 70% ethanol (2 times) at 5 minutes each. The slides were steamed for 35 minutes with a pH 6 Dako Target Retrieval (Agilent Technologies, S169984-2) and permeabilized with 0.1% Triton X-100 (Sigma-Aldrich, T8787) for 5 minutes, followed by a 5-minute wash with PBS. Slides were then blocked in serum-free, protein blocker (Agilent Technology, X090930-2). Slides were incubated with a biotinylated anti-pimonidazole mouse IgG1 monoclonal antibody diluted 1:50 in Background Sniper (Fisher Scientific, 5082385) for one hour at room temperature. This step was followed by washes with PBS. Streptavidin, AlexaFluor 568 conjugated secondary antibody (Life Technologies, S11226) was applied for 30 minutes at a 1:200 dilution in the dark at room temperature followed by washes with PBS. Slides were then mounted using ProLong Gold anti-fade with DAPI (Molecular Probes, P36935) for immunofluorescence staining. Images were acquired using a Leica fluorescence microscope keeping identical exposures across the samples, per channel. Quantitation of images was carried out via the Image J module, by measuring the percentage signal across the area of the field in each channel of interest.

### Statistical analyses

Specific descriptions of statistical analyses and experimental/biological replicates are provided in the figure legends and in relevant Methods subsections. Data are presented as ± standard error of the mean (SEM). Data were analyzed by two-tailed unpaired Student’s *t*-test with Welch’s correction, analysis of variance (ANOVA) for group analyses or correlation tests. Results with p-values < 0.05 were considered statistically significant, with * p< 0.05, ** p≤ 0.01, *** p≤ 0.001, **** p≤ 0.0001. Statistical analyses were performed using statistical software package R (v. 4.2.2) or GraphPad Prism (v. 9).

## ACKNOWLEDGEMENTS

This work was supported by R01CA254100 (PR), F31CA232653 (CT), R01CA254100-S1 CURE supplement (BMV), R01GM112415 (AB), a Sylvester Translational Science Grant (PR) and research support funds from the Sylvester Comprehensive Cancer Center (PR). JS was a scholar in the Sylvester K12 Calabresi clinical oncology research career development program, which is supported by the National Cancer Institute of the National Institutes of Health under award number K12CA226330. We thank Ms. Janiyah Vogle for assistance with manuscript preparation. We thank the following Sylvester Shared Resources: Bioinformatics and Biostatistics Shared Resource and Onco-Genomics Shared Resource. Research reported in this publication was supported by the National Cancer Institute of the National Institutes of Health under Award Number P30CA240139. The content is solely the responsibility of the authors and does not necessarily represent the official views of the National Institutes of Health.

